# Seeing the piles of the velvet bending under our finger sliding over a tactile stimulator improves the perception of the fabric

**DOI:** 10.1101/2024.03.22.586227

**Authors:** Laurence Mouchnino, Brigitte Camillieri, Jenny Faucheu, Mihaela Juganaru, Alix Moinon, Jean Blouin, Marie-Ange Bueno

## Abstract

Using friction modulation to simulate fabrics with a tactile stimulator (i.e. virtual surface) is not sufficient to render fabric touch and even more so for hairy fabrics. We hypothesized that seeing the pile of the velvet darken or lighten depending on changes in the finger movement direction on the virtual surface should improve the velvet fabric rendering. Participants actively rubbed a tactile device or a velvet fabric looking at a screen that showed a synthesized image of a velvet which either remained static (V-static) or darkening/lightening with the direction of touch (V-moving). We showed that in V-moving condition, the touched surface was always perceived rougher, which is a descriptor of a real velvet (*Experiment 1*). Using electroencephalography and sources localization analyses, we found greater theta band [5-7 Hz] oscillation power in the left inferior posterior parietal lobule (PPC) in the Virtual velvet/V-moving condition as compared to both Real velvet/ V-static and Virtual velvet/V-static conditions *(Experiment 2)*. This result is consistent with studies that give a crucial role to the left PPC for visuo-tactile binding. The greater activity of the lateral occipital area found in the Virtual velvet/V-moving condition could have contributed to the emergence of a velvet more realistic representation.

## 1. Introduction

Most of our daily interactions with the environment are based on goal-directed movements to explore and perceive materials or objects. This is mainly done by tactile exploration using the glabrous skin areas of the hand or foot. During this process, the brain actively modulates movement parameters, such as the contact force between the skin and the surface, the movement velocity and its direction. Meanwhile, the skin of the body segment in contact with the surface undergoes deformation during the motion thereby stimulating the tactile receptors. For example, microneurography recordings showed greater responses of both fast-adapting (FA) and slow-adapting (SA) tactile receptors (i.e., enhanced number and frequency of the spikes) when the direction of the movement of the finger in contact with hairy textile materials was against the main pile direction, as compared to along the pile direction (Breugnot et al., 2006). At first sight, what seems relevant for discriminating textures are the properties of the surfaces in contact (i.e., the skin of the hand and the materials) together with their relative motion. The friction-induced vibrations (i.e. FIV) that propagate in the skin during motion are also particularly relevant for the perception of fine textures (Faucheu et al., 2019; Manfredi et al., 2014; Massimiani et al., 2020). The vibration and the FIV would be mainly detected through Pacinian receptors (i.e. FAII), which are exquisitely sensitive to vibration (∼100-300 Hz, Johansson and Vallbo, 1983). However, finger-surface interactions do not only engage these receptors as FIV also stimulates fast- and slow-adapting receptors providing a rich and complex composition of tactile information (e.g., FAI, SAI and SAII, Dione et al., 2021).

Conceptually, the fact that fabrics discrimination strongly relies on friction modulation in a large frequency bandwidth from 0 to approximately 1000 Hz opens the possibility of creating devices capable of simulating different fabrics. Indeed, textiles have a minimal spatial period of some tens of microns and stimulators with friction modulation are good candidates to emulate fabrics (Bueno et al., 2014; 2015). The challenge is to give the illusion of touching a real fabric when exploring a tactile feedback device. Devices capable of reproducing surfaces have been designed by different groups using different technologies (e.g., Biet et al., 2008; Felicetti et al., 2022). Using ultrasonic vibrations to modulate the coefficient of friction between the finger and the surface, the STIMTAC device has been used in several tribology studies (Biet et al., 2008; Bueno et al., 2014; 2015; Camillieri et al., 2018). This device has proved to be interesting for reproducing textile fabric. Psychophysical investigations using the STIMTAC tactile device have shown that the perceptual rendering is promising (Weiland et al., 2024) but presents some limits in particular for hairy fabric (velvet) (Camillieri et al., 2018; Bueno et al., 2015). This perceptual discrepancy when touching real and simulated velvet could be partly due to the fact that the power of the respective FIV in the bandwidth [100–400 Hz] differed greatly (Camillieri et al., 2018). In addition, the response of the SAI units to the skin indentation (Merkel endings, Harrington and Merzenich, 1970) that should be produced when the finger is moving against the pile of the velvet, is lacking when contacting a tactile device.

Although the material qualities are mainly estimated by the sense of touch, the visual system is capable of providing rough information about material properties even before touching it (Tiest and Kappers, 2007; Baumgartner et al., 2013). For instance, the view of a surface with long curled hairs (like angora rabbit) is likely to be judged softer than a surface with short bristly hairs. Notably, the visual perception of the surface roughness is greatly dependent on the illumination direction: observers perceive surfaces to be markedly rougher with decreasing illuminant angle (Ho et al., 2006). Seeing how a surface is transformed when touching it can also provide information about the properties and identity of the surface. For instance, seeing dark and shinning traces during back and forth finger movements over the surface is likely to evoke a velvet-like fabric (a feature well exploited by painters such as Rembrandt who depicted the velvet fabric’s soft folds and luxurious sheen in the “Old man with beard, fur cap, and velvet cloak”). A possible mechanism underlying this phenomenon is that the view of the physical consequences of our interaction with these fabrics, would reactivate stored internal representations of the fabric properties that have been constructed during previous active explorations with the fabric (e.g., Romo et al., 2003; Zhou and Fuster, 2000).

The above-mentioned effects of visual information suggest that, during finger exploration, the appraisal of the surface properties might differ in contexts with and without surface-related visual feedback (see Driver and Spence, 2000 for a review). The various areas where tactile and visual sensory inputs converge in the brain could permit the visual inputs to influence tactile perception. This includes areas dedicated to the early (e.g., primary somatosensory cortex, Dionne et al., 2010; Zhou and Fuster, 2000) and subsequent stages of sensory processing (e.g., inferior premotor and inferior parietal cortex cortices; Sereno and Huang, 2014; Ishida et al., 2010). These brain areas contain cells that respond to either tactile or visual inputs, but also bimodal cells that respond to both tactile and visual stimulus. Internal representations and perceptual experience would be strongly contingent upon the activity of these regions. Hence, preventing the brain to be fueled with both visual and tactile inputs could limit the possibility of perceiving virtual fabrics through tactile devices.

In the present study, we tested the hypothesis that through sensory integration processes, immerging participants in a context that simulates finger friction/vibrations characterizing exploration of a velvet fabric, together with the visual (but virtual) consequence of the finger motion on velvet (i.e. shading/sheen changes) will improve the perception of the velvet. We used a twofold approach to test this hypothesis. First, the influence of the additional visual input on the perceptive attributes of simulated velvet fabric was analyzed through a behavioral study based on paired comparison tests *(Experiment 1).* We used the exploratory procedure (motor activity) for extracting particular object properties with accuracy and/or speed judgement (Klatzky et al., 1989). Specifically, this procedure necessitated local pressure of the fingertip during the exploratory movement to provide more detailed properties (e.g., softness, thickness, relief) as Giboreau et al. (2001). Our second approach is based on the current consensus that functional processing of sensory inputs is associated with the modulation of band-specific neural oscillation power (Fabre et al., 2023; Haegens et al., 2011; Pfurtscheller and Lopes da Silva, 1999). Notably, ongoing oscillations would shape our perception. In particular, oscillations within the low frequency bands (theta and alpha) would be linked to perceptual functions, notably to the perception or non-perception of a stimulus (Busch et al., 2009; Busch and VanRullen, 2010). We predicted that combining visual and tactile information when moving the index finger on a tactile device simulating the friction between the finger and a velvet fabric will modulate theta and alpha powers in the inferior parietal cortex (PPC), a key region for processing of tactile (Haegens et al., 2011) and visuo-proprioceptive information *(Experiment 2)*.

## 2. *Experiment 1*: Perception

### 2.1 Materials and Methods

Twenty-two participants (9 women and 13 men) who did not have any known neurological, physiological, cognitive or motor disorders participated to the experiment (20 right-handed and 2 left-handed, mean age: 27 ± 4 years). They all gave their written informed consent to take part in this experiment, which conformed to the ethical standards set out in the Declaration of Helsinki and which were approved by the CERSTAPS ethic committee (IRB00012476-2021-09-12-140). None of the participants had participated in any similar previous experiments.

The experiment took place in a room with conditioned atmosphere (20 ± 2°C and 65 ± 5% of relative humidity). Before each experiment, the participants washed and dried their hands. During the experiment, the participants were seated in front of a table on which the STIMTAC tactile device was positioned. A computer screen was located 40 cm behind the device. An adjustable support covered with a towel held the subject’s arm in a correct and comfortable position. The hand and the exploring surface were occluded from view by a custom-made box (Fig. 1). The STIMTAC is able to simulate different fabric-like materials (Ben Messaoud et al., 2016, Weiland et al. 2024) by inducing ultrasonic vibrations (at a constant value between 30-40 kHz) of the device surface relative to the finger position (Biet et al., 2008). Changing the amplitude of the vibration allows modulating the coefficient of friction (COF) between the finger and the surface, which is a key parameter for discriminating textures (Manfredi et al., 2014). With velvets (in particular pane velvet) the sensation of rubbing against the nap is very different to rubbing with the nap. To this end the STIMTAC coordinates the vibrations depending on the direction of movement to reinforce the sensation of a real velvet (Camilieri et al. 2018).

**Figure 1:**
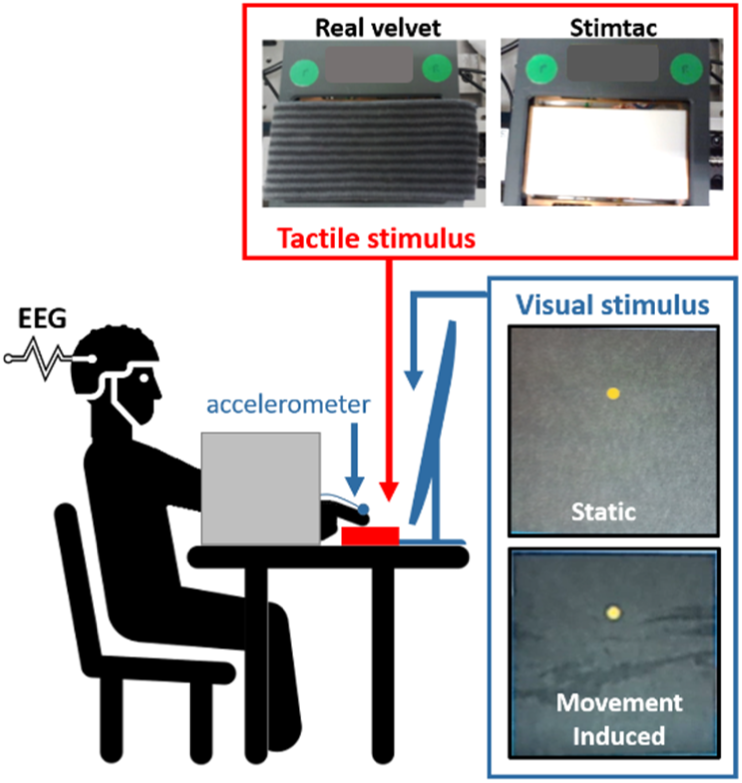
Experimental set up. The tactile stimulator STIMTAC is fixed on a three axes load cell. The screen in front of the participants shows either a static visual display (V-static) or a movement-induced visual feedback (V-moving). The right index finger is equipped with the accelerometer for the experiment 2.

In the present study, the STIMTAC generated two different tactile stimuli to simulate two different percepts. The T_Vel_ tactile stimulus reproduced the friction recording during exploration of velvet (as in Camillieri et al., 2018) while the T_Sham_ tactile stimulus is generated from T_Vel_ but with a lower magnitude, corresponding to no known fabrics. More specifically, the vibration amplitude used to generate the T_Vel_ stimulus was set at 74% of the STIMTAC’s maximum vibration amplitude, while the vibration amplitude of the T_Sham_ stimulus was set at 0.57*T_Vel_. Note that the measured mean perception threshold of the vibrations across participants was 0.19*T_Vel_.

The computer screen presented a 12 x 10 cm synthesized image of grey velvet with a yellow dot on which the participants fixated throughout the trials (see Fig. 1). During the tactile exploration of the surface (see below), the visual display either remained static (V-static) or simulated the traces that would have been left if the participant’s index finger was sliding on a velvet fabric (V-moving). The STIMTAC device records the position of the finger enabling the virtual velvet image to be modified in real time. More specifically, when the participants rubbed the surface against the nap (or grain)of the simulated pile (i.e. towards the right in this protocol), the visual display left a dark trail (as if the velvet fibers straightened). When they rubbed the surface along the nap of the simulated pile (i.e. towards the leftward direction), the traces on the screen became lighter (as if the velvet fibers flattened). The spatiotemporal characteristics of the traces displayed on the screen matched those of the index finger movements on the STIMTAC surface (a crucial aspect for visuo-somatosensory integration, see Hidaka et al., 2015 for a review).

The participants were asked to produce back and forth lateral movements with the index finger of their right hand to explore the stimuli programmed on the STIMTAC tactile stimulator. The mean amplitude and velocity of the sliding finger movements were ∼40 mm and ∼40 mm/s, respectively. A 1 Hz metronome beat helped the participants to achieve the desired movement velocity. The instructions specified that the normal force on the surface had to be ∼0.5 N. Before the experimental session, the participants were trained to produce finger movements which complied with all these specifications. For this training session, two dots, 40 mm apart were positioned just in front of the exploring surface and a monitor provided the normal force feedback to the participants (see Fig. 1). During the training session, the finger movements were performed on a different fabric. Only a few back and forth movements were necessary for the participants to comply with the required movement features.

For the experiment, four different scenarios were designed according to the selected combination of tactile (i.e., T_Vel_ or T_Sham_) and visual stimuli (i.e., V-static or V-moving) (see Table 1).

**Table 1:** Scenarios used in Experiment 1.

| STIMTAC<br>Command signal | Visual feedback |  |
| --- | --- | --- |
|  | V-moving | V-static |
| $T_{Vel}$ = virtual velvet | *Scenario 1* | *Scenario 2* |
| $T_{Sham} = 0.57 * T_{Vel}$ | *Scenario 3* | *Scenario 4* |

The task of the participants was to compare the roughness between the surfaces presented in two distinct sets of scenarios. For each paired comparison test, the participants indicated “which surface is rougher”? The participants were instructed that they could answer that the two surfaces had the same roughness. The French term “râpeux” was selected for “rougher” among several French terms describing rough textures from Bassereau and Charvet-Pellot (2011). We chose this descriptor to describe the effect of the pile tuft under the fingers which can well characterize the “velvet effect”, i.e. the difference between the feeling when moving the finger along and against the pile direction. At the beginning of the test, the participants explored the first presented scenario before exploring the second scenario. They were allowed to explore each surface for as long and as many times as required. The order of presentation of the different scenarios was based on a Latin square-like design allowing for 16 pairs of scenarios randomly presented for each participant. Therefore, each scenario was presented as often in first and second positions. For the analyses, the pairs were categorized according to the sequence of the scenario presentation (e.g., 1vs4, 4vs1, 4vs2, …). Each participant touched a sample of real velvet before starting the experiment, and at three moments equally spaced in time during the experiment to refresh the “velvet effect”.

### 2.2. Statistical analyses

In *Experiment* 1, for each paired comparison, based on the votes provided by all 22 participants, the frequency of occurrence of one scenario being perceived rougher is observed for each pair of scenarios.

### 2.3. Results

The overall analysis of the collected data showed evidence of a position bias which is commonly observed in paired comparison experiments (Penny et al., 1972; Day, 1969). The position bias was observed when considering all paired comparison results in that the vote count of all 22 participants showed that the tactile stimulus for the scenario presented in the second position was more often perceived rougher than for the scenario presented in the first position (161 vs 110 votes). The position bias was strong for (2vs2), (3vs3) and (4vs4) tests, indicating that the tactile stimulus of the scenario presented in second position was incorrectly perceived rougher than for the same scenario presented in first position. Irrespectively of the tactile stimulation contents, the second scenario was perceived more frequently by the participants as being rougher (f_2_=0.458) than the first scenario (f_1_=0.312). The effect of the scenario position was considered significant as the confidence interval was small: [f-0.05; f+0.05]. Importantly, for the 1vs1 tests, the scenarios were mostly perceived (correctly) as the same scenario which tended to indicate that *Scenario 1* (i.e. Virtual velvet with the V-moving) was more clearly perceived than the three other scenarios.

To take into account the position bias, the collected data were analyzed with regards to the order of presentation. The analysis focused on the reference set of scenarios, in particular for pairs including *Scenario 1 (Virtual tactile velvet / V-moving)*. The observed frequencies of tactile stimulus of *Scenario 1* being perceived rougher than in another scenario when presented in either first or second position was calculated (Table 2). Relative to *Scenarios 3 and 4*, tactile stimulus in *Scenario 1* was perceived rougher when presented in both first and second position. The result was consistent with the fact that the tactile stimulus in Scenario *1* reproduced the FIV recorded when exploring real velvet (i.e., rough fabric) while in *Scenarios 3 and 4,* the tactile stimulus reproduced only a fraction of the original signal (i.e., 57%). More importantly, the tactile stimulus in *Scenario 1* relative to *Scenario 2* was also perceived rougher when presented in both first and second position. Remarkably, in this case, the vibration characteristics of *Scenarios 1 and 2* were exactly the same. The only difference between these Scenarios was in the type of visual feedback associated with the explorative finger movements. *Scenario 1* was associated with V-moving while *Scenario 2* was associated with V-static. This finding indicated that the movement-induced visual feedback sharpened the roughness perception. It also suggested that, alone, the movement-induced visual feedback provided during finger exploration had limited capacity to create the illusion that one was touching real velvet. Indeed, in comparison with S*cenario 1*, tactile stimulus in *Scenario 2* was perceived rougher than in *Scenario 3*, and the observed frequency of occurrence of the tactile stimulus in *Scenario 3* perceived rougher than in *Scenario 4* was not higher. This indicated that the sharpening effect of the movement-induced visual feedback depends on the characteristics of the tactile stimulus.

**Table 2:**
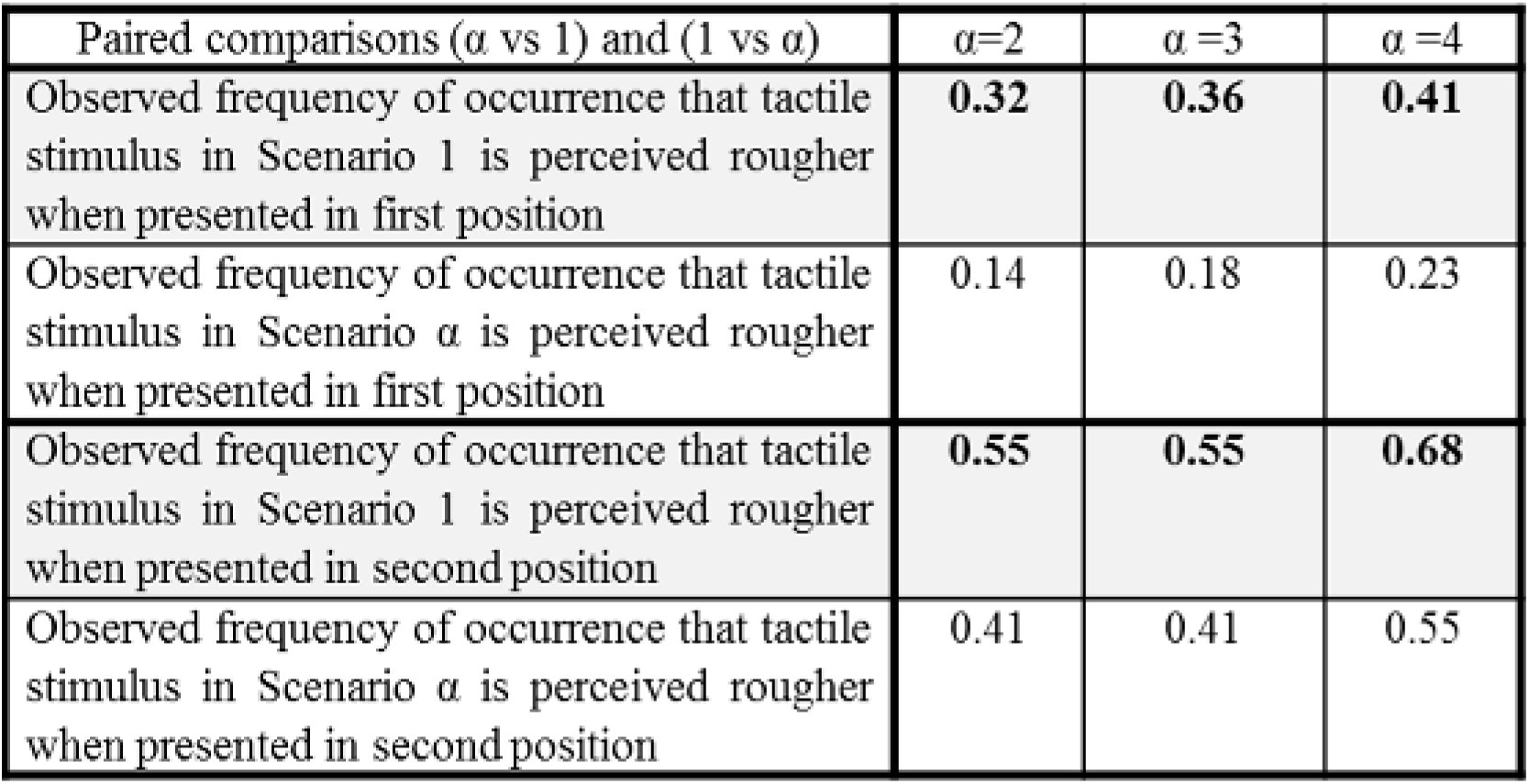
Observed frequencies that tactile stimulus in Scenario 1 (i.e. virtual velvet with V- moving) is perceived rougher than in the other scenarios.

| Paired comparisons ($\alpha$ vs 1) and (1 vs $\alpha$) | $\alpha=2$ | $\alpha=3$ | $\alpha=4$ |
| --- | --- | --- | --- |
| Observed frequency of occurrence that tactile stimulus in Scenario 1 is perceived rougher when presented in first position | **0.32** | **0.36** | **0.41** |
| Observed frequency of occurrence that tactile stimulus in Scenario $\alpha$ is perceived rougher when presented in first position | 0.14 | 0.18 | 0.23 |
| Observed frequency of occurrence that tactile stimulus in Scenario 1 is perceived rougher when presented in second position | **0.55** | **0.55** | **0.68** |
| Observed frequency of occurrence that tactile stimulus in Scenario $\alpha$ is perceived rougher when presented in second position | 0.41 | 0.41 | 0.55 |

## 3. *Experiment* 2: Tribological measurement and electroencephalography

### 3.1 Materials and Methods

The experiment was carried out with the same ethical and environmental conditions as in *Experiment* 1. Fourteen right-handed participants (10 women and 4 men, mean age: 27 ± 2 years) without any known neurological, physiological, cognitive and motor disorders participated in the experiment. None had participated in any previous similar experiments. The Semmes–Weinstein monofilament test was carried out to determine the sensibility of the index finger pad of each participant. The sensitivity thresholds varied between 0.023 g to 0.166 g (mean 0.11 grams ± 0.06) and were therefore deemed as ‘normal’ (Hage et al., 1995).

The same set-up as in *Experiment* 1 was used in *Experiment* 2 (e.g., computer screen, STIMTAC device) with the addition of an EEG system to record cortical activities. At the start of each trial, the participants had to put their right index finger pad on the left extremity of the virtual (STIMTAC) or real surface and to maintain gaze on the dot on the screen. They were instructed to stay relaxed and to focus on the material they were touching with their finger. The participants were asked to explore the surface with back and forth lateral movements of the finger continuously during 24 s. The speed (∼40 mm/s) and the amplitude (∼40 mm) of the finger movements as well as the exerted normal force on the surface (i.e., ∼0.5 N, Fig 2), were similar to those produced by the participants of *Experiment* 1.

**Figure 2:**
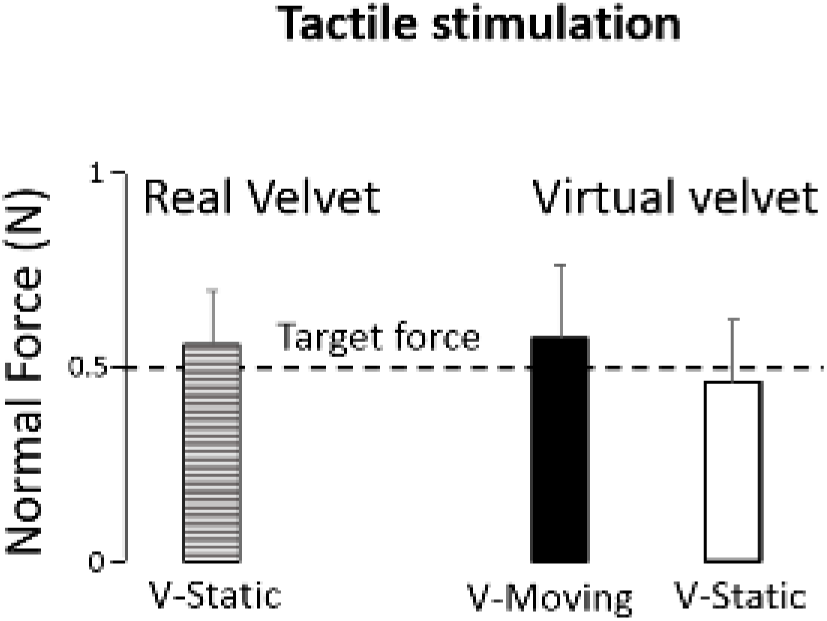
Mean normal force for all participants (N = 14). Error bars represent standard deviation across participants.

Two surfaces from *Experiment* 1 were tested: the real velvet fabric and the surface (STIMTAC device) that simulated the velvet friction (hereafter referred to as virtual velvet). For trials with the real velvet, the synthesized image of grey velvet remained static (V-static). For trials with the virtual velvet, this image provided either a movement-induced visual feedback (V-moving) or remained static (V-static) as in *Experiment* 1 (see *Scenarios* 1 and 2). For trials with V- moving, the visual display was refreshed at the end of the trial (i.e., it appeared as in V-static display conditions). Stationary trials (no finger movement with V-static) were also recorded during 24 s and served to analyze EEG signals (see below). For reasons of homogeneity, the same auditory cues (i.e. metronome beats) as in trials with surface exploration were also delivered for the stationary trials.

For each experimental condition, the participants performed 4 trials to ensure 40 back and forth movements (i.e. Real velvet with V-static, Virtual velvet with V-static and Virtual velvet with V-moving). Four stationary trials were also recorded. The order of the condition presentation was randomized within the experiment and across participants.

#### Tribological measurement: Finger friction and induced vibrations

The real fabric or the STIMTAC were affixed directly on a 3 axes load cell (model 3A60-20 N, Interface Inc., Scottsdale, Arizona, recording at 2000 Hz). It provided the components of the force exerted by the finger along three orthogonal axes (the normal force F_n_ is along the vertical axis (perpendicular to the surface); the axes in the horizontal plane allow the calculation of the tangential force F_t_). From these two forces the instantaneous coefficient of friction (COF), corresponded to F_t_ / F_n_, is calculated. We measured both F_t_ and F_n_ and, the finger vertical acceleration while the participants were exploring the surfaces with their index finger. To measure the finger vertical vibrations induced by the friction, an accelerometer (Piezoelectric Charge Accelerometer 4374 with a charge amplifier 2635 from Bruël & Kjaer, Mennecy, France) was glued on the skin just above the nail of the right index finger. The data were acquired with Pulse software (Bruël & Kjaer, Mennecy, France). The accelerometer signal (recorded at 2000 Hz) was analyzed in the frequency domain. The variables extracted from the vibration signal are the root mean square of the acceleration (RMS) and the spectral power for 4 frequency bandwidths [3-100 Hz], [100-200 Hz], [200-400 Hz] and [400-800 Hz], following the methods and results previously described in Camillieri et al. (2018).

#### Brain activity (EEG)

EEG activity was recorded continuously using a Biosemi ActiveTwo system (The Netherlands, 64 Ag/AgCl electrodes, 1024 Hz sampling frequency). The EEG data were pre-processed using BrainVision Analyzer2 software (Brain Products, Gilching, Germany). EEG signals were referenced against the average of the activities recorded by all electrodes. Then, 2000 ms long epochs were extracted from the EEG signals and synchronized with respect to the point in time at which the participants started to move their index finger against the (real or simulated) velvet piles (total of 40 epochs for each participant). The onset of the movement was defined as the initial change in the tangential force towards the right (i.e., against pile). Epochs were visually inspected and those presenting artifacts were rejected. On average (across the participants), 38-40 epochs were included in the analyses.

EEG neural sources were estimated with the Dynamical Statistical Parametric Mapping (dSPM, Dale et al., 2000) implemented in the Brainstorm software (Tadel et al. 2011), which is documented and freely available for download online under the GNU general public license (http://neuroimage.usc.edu/brainstorm). A boundary element model (BEM) with three realistic layers (scalp, inner skull and outer skull) was used to compute the forward model on the anatomical MRI brain template from the Montreal Neurological Institute (MNI Colin27). Using a realistic model provides more accurate solution than a simple three concentric spheres model (Sohrabpour et al., 2015). We downsampled the cortex surface to 15 002 vertices which allowed good spatial resolution. Measuring and modelling the noise contaminating the data is beneficial to source estimation. Noise covariance matrices were computed using the stationary trials (i.e. while the participants remained still). Such EEG source reconstruction has proved to be suited for investigating the activity (which is indexed to current amplitude, Tadel et al., 2011) of outer and inner cortical surfaces with 64 sensors (Ponz et al., 2014). The current maps were averaged over 4 time-windows of 500 ms from the start of the shear forces in the against pile main direction to the end of the along pile movement direction for each participant, surfaces and visual conditions. Note that due to the metronome beats participants were accurate in completing each one-way rubbing movement in 1000 ms.

Time frequency analyses were performed in EEG source space. The data were transformed into time-frequency domain using Morlet wavelet transforms. The respective power of theta (5-7 Hz) and alpha (8-12 Hz) cortical oscillations were computed for each trial and then averaged across trials for each condition and participant. We normalized the frequency powers by computing averaged theta and alpha event-related synchronization / desynchronization (ERS/ERD, 2000 ms epochs taken for the static trials). These computations were performed in regions of interest (ROIs) from the angular gyri of the right and left posterior parietal cortices (PPC, Brodmann area 39). These ROIs were manually defined, based on the Destrieux cortical atlas (Destrieux et al., 2010), and had both 242 vertices. We purposely selected two time windows, one from 500 to 1000 ms and one from 1000 to 1500 ms to compare changes in brain electrocortical activities (ERS/ERD) between movements against (500 to 1000 ms) and along (1000 to 1500 ms) the main pile direction. These two time windows allowed the removal of any edge effects as wavelet coefficients are less accurate at the beginning (here 0 ms) and end (here 2000 ms) of a time series (Torrence and Compo, 1998).

### 3.2. Statistical analyses

The analyses of the finger coefficient of friction and FIV were performed with the XLSTAT software. The objective was to compare, for each variable, two sets of data (e.g., Real velvet – V static vs Virtual velvet – V static) to determine if they belonged to the same population or not. In all cases, the data from the two sets were obtained from the same participants (paired data). Firstly, we verified whether each set of data followed the normal law with the Shapiro-Wilk test (p-value > 0.05). If the sets followed the normal law, the variances were compared with the Fisher-Snedecor test (F-test). In all cases for which the variances were not significantly different, the means were compared with the Student’s t-test for paired data. If one or both of the two sets did not verify the normal law, the Wilcoxon signed rank test for paired data was used.

One-way ANOVAs were used to assess the effect of Condition (i.e., Real velvet with V-static, Virtual velvet with either V-static or V-moving) on the normal force, and on the theta and alpha band powers (separate analyses for the left and right inferior PPC). The time frequency analyses were performed separately for the left and right inferior PPC. For these analyses, the theta and alpha band mean powers, significant effects (statistical threshold of p < 0.05) were further analyzed using Newman-Keuls post-hoc tests. All data had normal distributions (as confirmed by Kolmogorov-Smirnov tests). We used t-tests (p < 0.05, FDR correction for multiple comparisons) to contrast the cortical current maps computed when the participants explored the real velvet and when they explored the virtual velvet (in either the V-static or V-moving visual display).

### 3.3. Results

#### Tribological results

The participants complied with the requirement to exert a ∼0.5 N normal force with the index finger during the surface exploration (overall mean of 0.51 N ± 0.15, Fig. 2). The ANOVA showed that this force did not significantly differ between the different visual and tactile conditions (F_2,26_ = 2.30; p = 0.11).

Figure 3A shows for each participant that the COF computed when exploring the Virtual velvet (ranging from 0.1 to 1.5) was generally much higher than the COF computed when exploring the Real velvet condition (ranging from 0.3 to 0.9). The linear regression equation estimated from these data attests that, for most of the participants, the COF was greater for the Virtual velvet than for the Real velvet rubbing movements. Figure 3B shows that for the same Virtual velvet surfaces, there is no influence of the visual display on the COF. This result was expected as the COF mainly depends on the skin/surface interaction.

**Figure 3:**
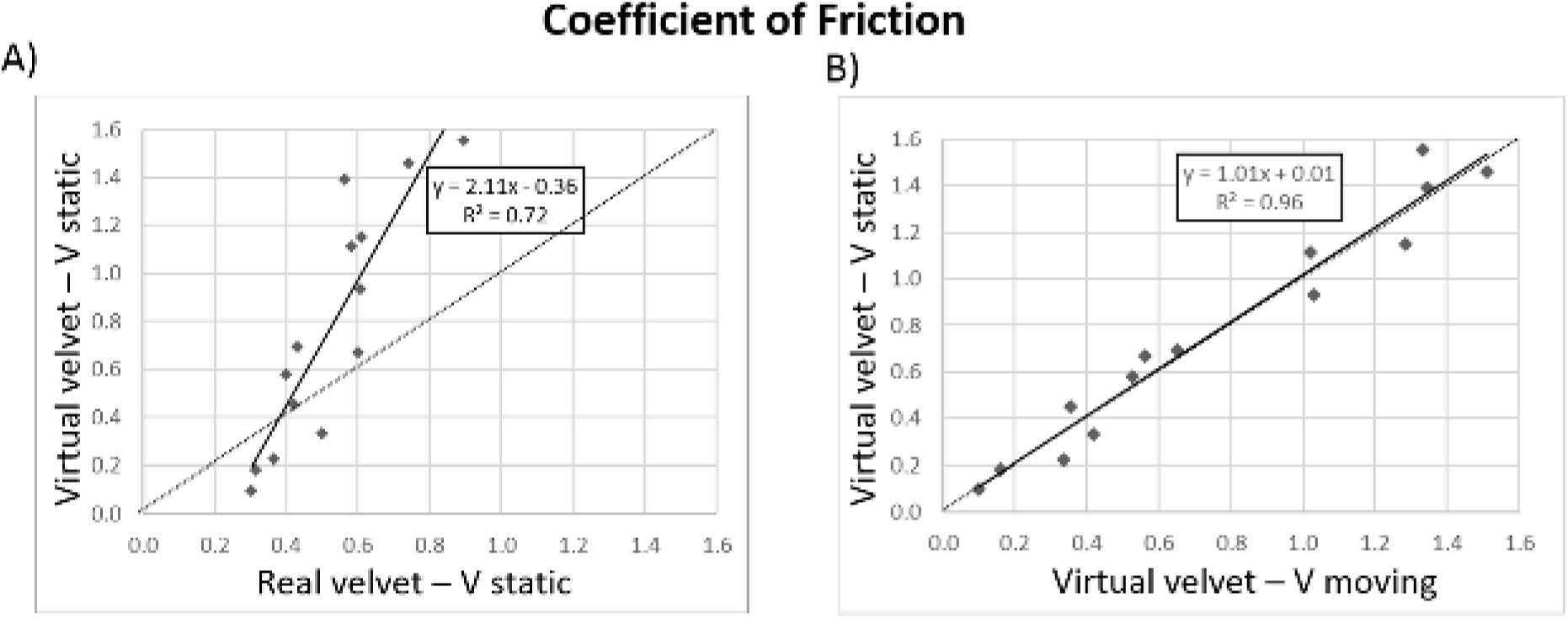
Coefficients of friction for all the participants: **A)** for virtual and real velvet fabrics (the dashed line is the line y = x), **B)** for the virtual velvet for V-static and V-moving visual feedback. A dot corresponds to a participant.

Figure 4 shows the FIV (i.e. the RMS of the finger vertical acceleration) for the 3 surface/visual display conditions (Fig 4A-C). The averaged FIV autospectra had the same global shape between conditions. However, across the FIV frequencies, the RMS distribution was smaller when the participants explored the real velvet compared to the virtual velvet. Moreover, in conditions with a V-static display, exploring the virtual surface yielded greater RMS than when exploring a real velvet with static display (Fig. 4D). For the virtual velvet, the autospectra was very similar between both visual stimuli (i.e., V- static and V-moving, Fig. 4E).

**Figure 4:**
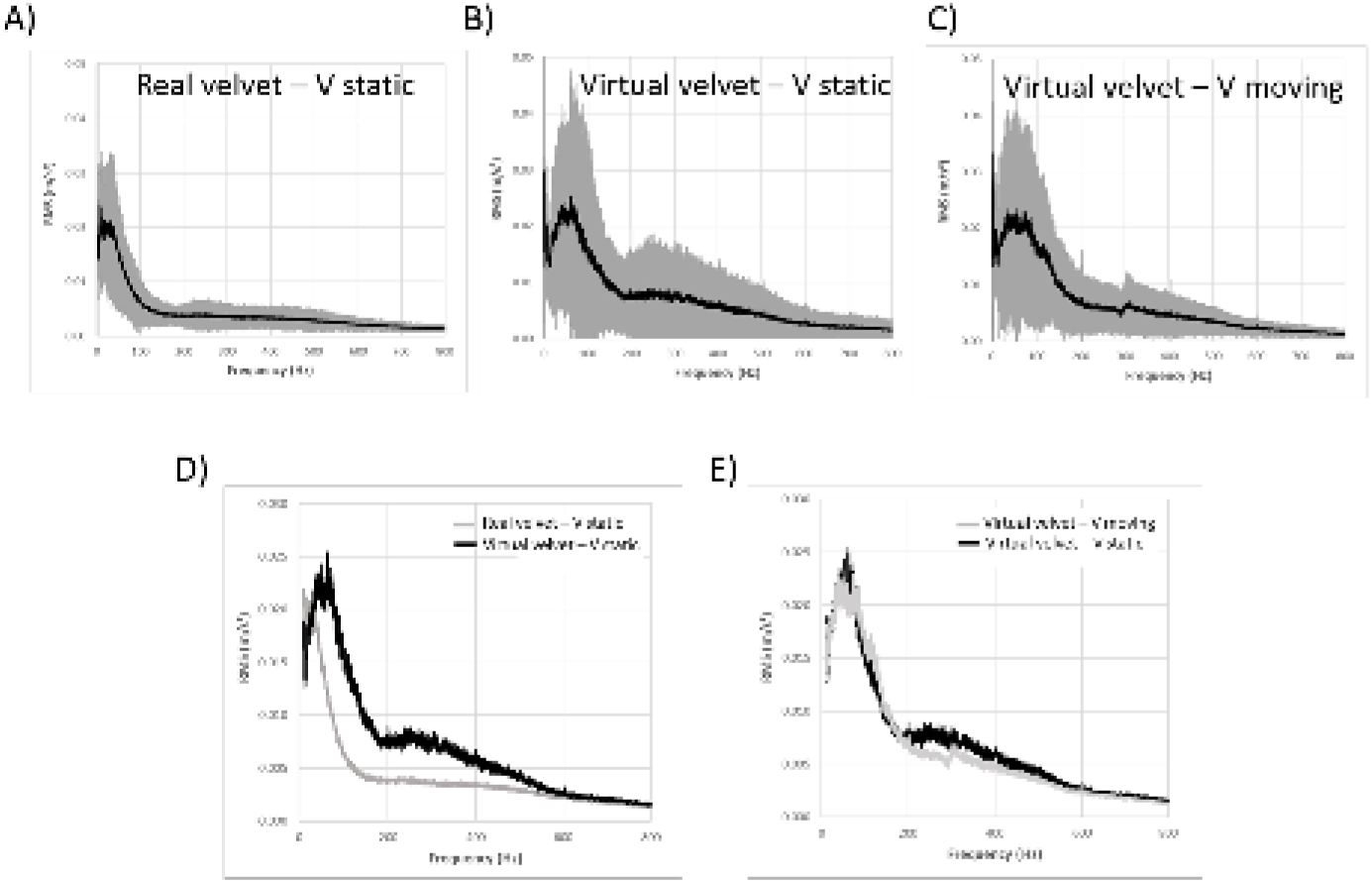
Acceleration autospectra obtained for all the participants. The average autospectrum is in black and the standard deviation in grey. **A)** for the real velvet with V-static visual display, **B**) for the virtual velvet with V-static and **C**) for the virtual velvet with V-moving. **D**) Average acceleration autospectra for all the participants obtained for the real and virtual velvet with V- static visual display, **E**) for the virtual velvet with V-static or V-moving.

The box plots of the spectral power (Fig. 5) confirmed that the variability between the participants was lower with the real velvet, whatever the frequency bandwidth. Moreover, the p-values for the spectral powers obtained for the different bandwidths show that the FIV when rubbing the real velvet and the virtual velvet with V-static were significantly different between 100 and 400 Hz (ps < 0.05, Fig. 5B-C). In addition, with virtual velvet, the type of visual stimulation had no significant influence on the FIV (ps>0.05, Fig. 5A-D).

**Figure 5:**
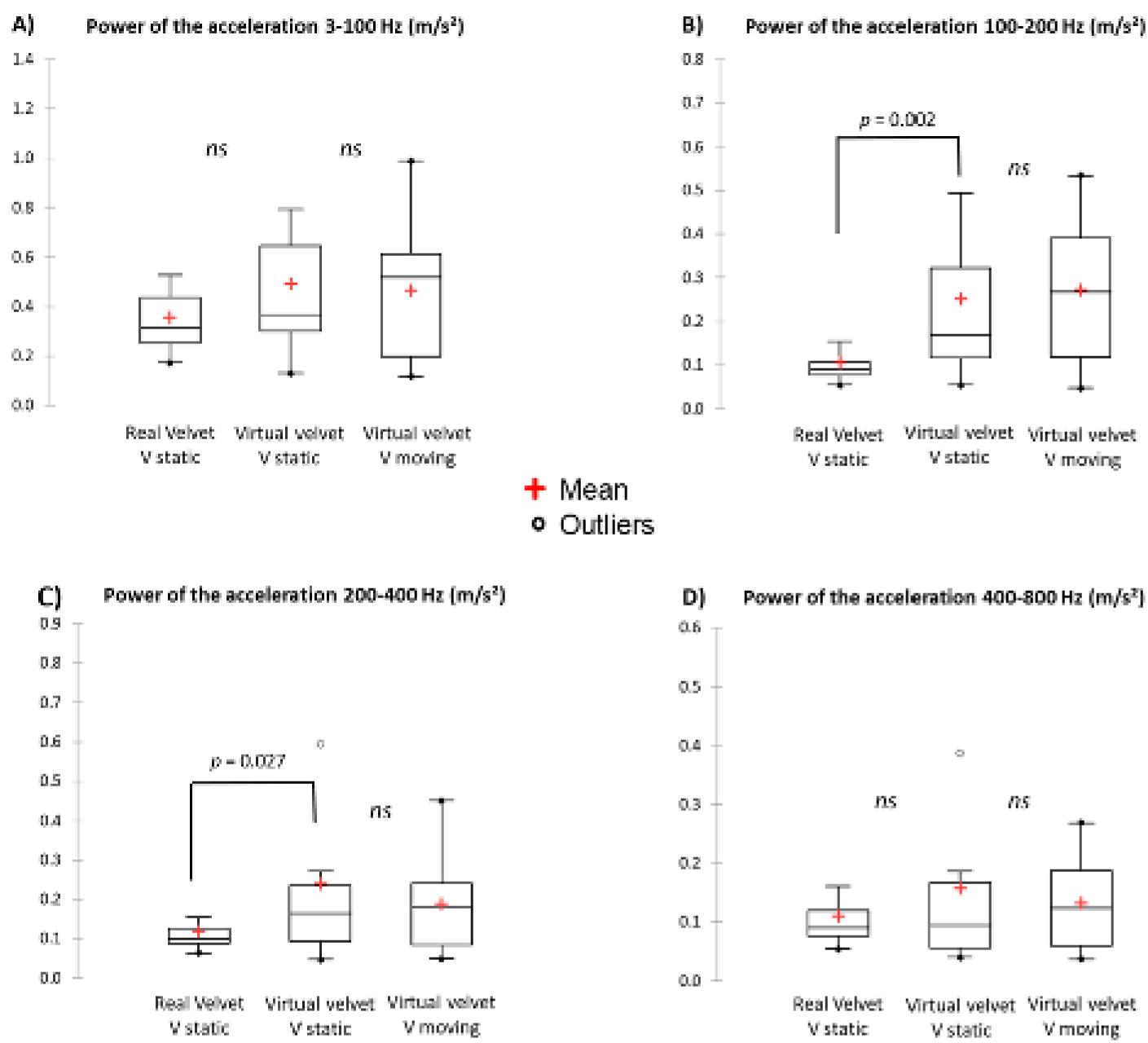
Box plot for the spectral power of acceleration for all the participants obtained for the real velvet with V-static, for virtual velvet with V-static and V-moving for different bandwidths: **A**) 3 to 100 Hz, **B**) 100 to 200 Hz, **C)** 200 to 400 Hz and **D**) 400 to 800 Hz. The middle line in the box plot is the median and the red cross the mean.

#### Brain activity

##### Surface-specific source localization

Figure 6 shows the statistical cortical maps for two different contrasts. As a striking result, the topography of the cortical activations estimated by source analyses differed greatly between the V-moving and V-static (compare Fig. 6A and Fig. 6B). The effect of the visual display was particularly noticeable when contrasting the real with the virtual velvet with V-moving (attested by the cold/blue colors of the cortical regions in Fig. 6A). Indeed, during virtual velvet finger exploration, the lateral occipital (LOC) and inferior PPC, and the medial surface of the left PPC (precuneus) of the right hemisphere showed significantly greater activation in the contrast Real velvet/V-Static minus Virtual velvet/V-moving conditions but not in the contrast Real velvet/V- Static minus Virtual velvet/V-static. The greater activity of the PPC observed with the Virtual surface persisted throughout the exploring finger movements, except for the left precuneus whose activity was significantly greater only at the beginning of the movement against pile main direction [0-500 ms]. Note that in both contrasts (Fig. 6A-B), the right dorsal anterior cingulate cortex (ACC) showed greater activation when moving on the virtual surface solely at the beginning of the movement against pile [0-500 ms] regardless of the visual feedback.

**Figure 6:**
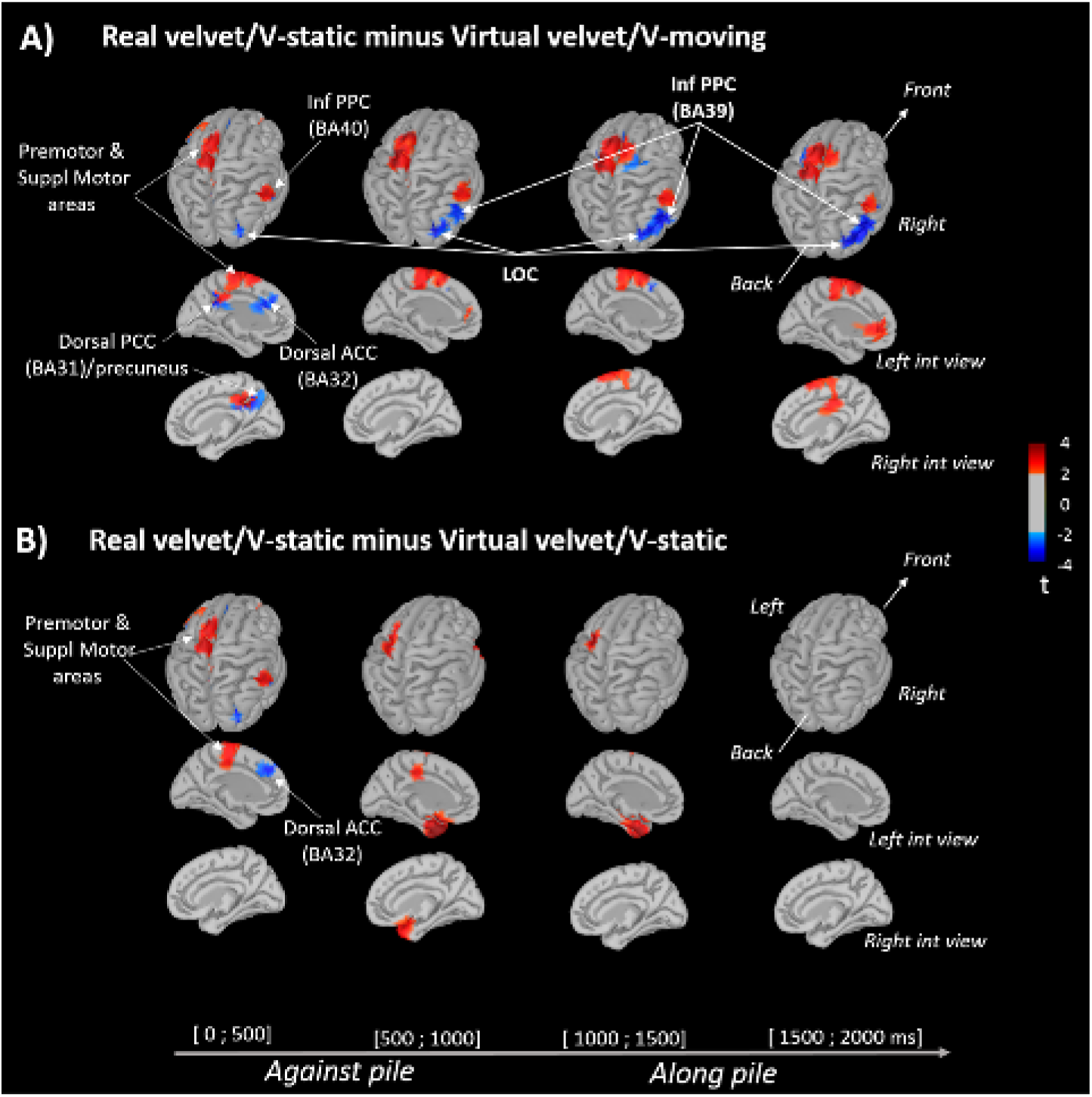
Statistical source estimation maps for real velvet versus virtual velvet fabric with V- moving (**A**), and versus virtual velvet with V-static (**B**) contrasts. Significant t-values (p ≤ 0.05, n = 14) of the source localization were shown during the 4 time window of 500 ms starting at the movement against pile main direction onset to the end of the along pile movement. Sources are projected on a cortical template (MNI’s Colin 27). For the first 500 ms contrast, we display the top and the left inner cortical views.

Moreover, moving the finger on a real velvet fabric engaged greater activity of the left motor area (i.e., contralateral to the moving finger) than when moving on a virtual velvet (i.e. warm/red colors in pre-rolandic cortex, Fig. 5). Interestingly, this enhanced motor activity was observed when the participants moved their index finger both against and along the pile in the V-moving condition, but only when the finger moved against the pile in the V-static display condition. The increased activation of the left sensorimotor region with real velvet was accompanied with a greater activity of the right PPC BA 40 (rostral to BA 39), that was only observed when participants received V-moving feedback during their movement.

#### Modulation of theta (5-7 Hz) oscillations in the left inferior PPC

Time-frequency analyses performed on the left and right PPC ROI (Fig. 7A) showed that theta band power was significantly impacted by the conditions (surface/visual feedback). This effect was observed for the left PPC when the participants moved their index finger along the pile (F_2,24_ = 5.55; p = 0.01, Fig. 7B, right panel). Post-hoc analyses revealed that the mean theta power was greater in the Virtual velvet/V-moving condition than in both the Real velvet/V- static (p = 0.013) and Virtual velvet/V-static (p = 0.015) conditions. Note that a significant desynchronization (ERD) was observed for both conditions with a V-static visual display (t-test for means against a value 0; t_12_ = -3.36, p = 0.0055 and t_12_ = -3.59; p = 0.0036, respectively for the Real velvet and Virtual velvet). No significant effect of condition was observed on theta band power for the movements against pile (Fig. 7B, left panel F_2,24_ = 2.79; p = 0.08) or for the right PPC (F_2,24_ = 1.04; p = 0.36 and F_2,24_ = 1.47; p = 0.24 for the along pile and against pile movement, respectively). The effect of condition on the alpha band power computed in the left and right PPC ROIs was not significantly different (ps > 0.05).

**Figure 7:**
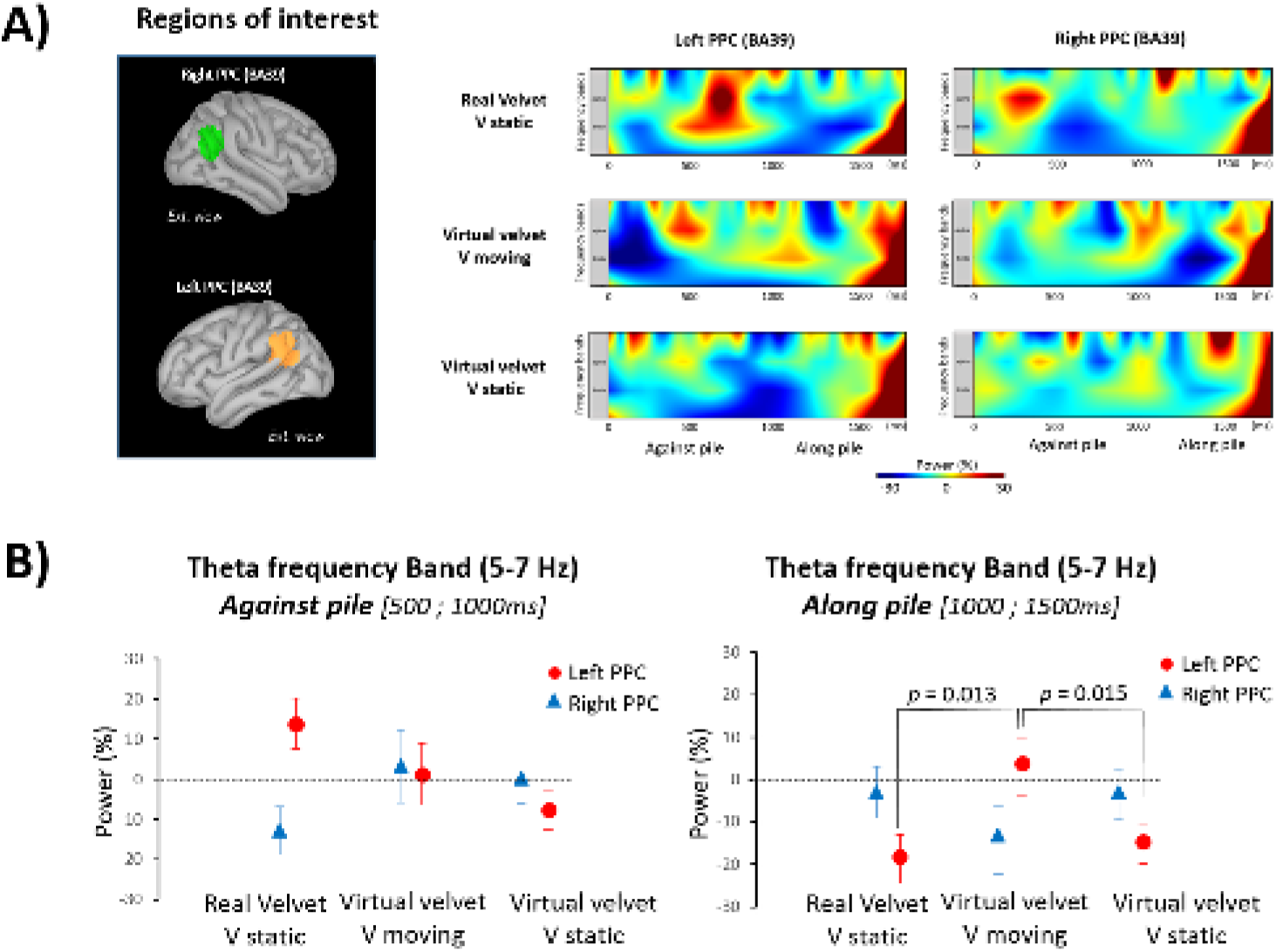
**A**) Time-frequency power (ERS/ERD) of the signals by means of a complex Morlet’s wavelet transform applied on the ROIs for each trial of each participant then averaged across participants. Cooler colors indicate ERD (event-related desynchronization) while warmer colors indicate ERS (event-related synchronization). Frequency bands of theta and alpha bands are illustrated to present changes in brain electrocortical activity. **B)** Group means power for Theta (5-7 Hz) frequency band computed during [500; 1000 ms] (left panel) and [1000; 1500 ms] (right panel) time windows (error bars depict standard error of the mean) over the left and right PPC.

## 4. Discussion

We investigated the perceptual rendering of virtual velvet fabric from tactile simulation (*Experiment 1*) together with the behavioral and neurophysiological substrates subserving this perception (*Experiment 2*). Our core objective was to test the possibility of improving velvet fabric rendering of a tactile device by adding the visual effects of rubbing a velvet fabric with a finger. The originality of our study lies in the fact that the participants could not see their moving finger (vision of the hand was occluded), but only a visual simulation of the trails left on a velvet fabric Specifically, when the participants moved their index finger against or along the main direction of the pile simulated by the tactile device, the visual display left a darker and lighter trace, respectively.

In a previous experiment, it was shown that the STIMTAC device used in the present study failed to simulate real velvet fabric (Camillieri et al., 2018). Our results suggest this could be due, in part, from the COF and the RMS of the vertical finger acceleration which differed when our participants explored the virtual velvet and the real velvet (*Experiment* 2). Remarkably, however, when the same tactile stimulation was combined with the visual rendering of a real velvet being rubbed by the participant’s finger, the surface of the explored tactile device was perceived as being rougher (*Experiment* 1). Because the attribute “rough” is a predominant velvet descriptor (Bassereau and Charvet-Pellot, 2011), this result could be construed as evidence that the dynamic velvet-like visual simulation enhanced the tactile-induced velvet perception. This enhancement may be due to the increase in roughness related to the against the grain movement.

Our perceptive and behavioral results then point to a crossmodal integration between tactile and visual information that can alter the touch sensation. Note, however, that the “velvet effect” (i.e., descriptor chosen to describe the effect of the pile tuft under the fingers) was not observed when the dynamic velvet image (V-moving) was combined with a tactile stimulus that did not relate to any known fabrics (i.e., sham stimulation in *Experiment* 1). This might suggest a near-effective skin/surface interaction in the present T_Vel_ condition for enabling a velvet rendering. This hypothesis is supported by studies showing that near sensory threshold stimulations can be perceived if combined with coherent stimulation from other sensory modalities (e.g., Popov et al., 1999; Dalton et al., 2000). It is also in line with those studies showing a more efficient integration of visual feedback in conditions with degraded somatosensory inputs (e.g., Mizelle et al., 2016; Tsay et al., 2021, see Limanowski, 2022 for a review).

The increased activity observed in the occipital and inferior parietal lobes (*Experiment 2*) could underlie the crossmodal sensory processes when velvet-like movement-induced visual feedback and tactile stimulus were combined. This would be consistent with the fMRI study of Limanowski and Blankenburg (2017) showing that these two posterior cortical regions work together to evaluate visuo-tactile congruence between the seen and the felt (tactile) hand positions. Increasing the congruence between visual and tactile cues during the active finger movements may have prompted the binding effect that denotes the mutual attraction between the visual perception of the velvet pile bending under the caressing finger and the simulated tactile inputs generated by the sliding finger on the tactile device.

In the left PPC, the power of theta band oscillations was markedly greater in the condition combining velvet-like tactile stimulation and movement-induced visual feedback than in both conditions with a static visual display (with either real or virtual velvet tactile stimulation). Interestingly, in the former condition, the participants of *Experiment 1* perceived the explored surface as being rougher than in both conditions with a static visual display, which used velvet-like or sham tactile stimulations). Previous studies showed evidence that increases in theta power enhance visuo-tactile integration processes (see Kanayama & Ohira 2009). In the present study, the greater theta power was observed over the left PPC, i.e. in the hemisphere contralateral to the moving finger, when the participants’ index finger was moving along the pile. Because integrative processes take ∼250 ms (see Kanayama & Ohira 2009), the fact that the increased theta power was observed from the very beginning of the finger movement along the pile direction suggests that this change of theta power started during the against pile /along pile reverse movement. Theta oscillations could be instrumental in binding visuo-tactile information to increase touch sensation. This is supported by studies showing that higher amplitudes of theta oscillations encode for touch intensity (Michail et al., 2016), fabric physical properties (e.g., warmness, softness, overall comfort, Jiao et al., 2020) as well as for more salient sensory stimuli (Iannetti et al., 2008). The dynamic velvet-like visual feedback could have also primed tactile representation of the velvet fabric. Such priming effect is consistent with the Brunyé et al.’s (2012) discovery that reading about tactile properties affects the subsequent tactile perception. More specifically, the authors found that all fabric ratings became smoother after reading a sentence implying a smooth tactile property, and rougher after reading a sentence implying a rough tactile property. In *Experiment* 1, the participants perceived the virtual tactile surface as being roughest in the condition that combined the velvet-like tactile, i.e. T_Vel_, and dynamic visual stimulations. Combining either movement-induced visual feedback with sham tactile stimulation or velvet-like tactile stimulation with a static image of a velvet fabric decreased the roughness perception of the explored surface. Together, these findings suggest that visuo-tactile sensory integration and sensory priming effect are not the most critical aspects for perceiving a tactile device as being rough. Rather, the relatedness between tactile and visual feedback clearly emerged in the present study as the most relevant factor.

The greater activations of the right occipital and lateral occipital cortices and of the inferior PPC (BA 39) were found in the condition with velvet-like dynamic tactile and visual stimulations compared to the condition with real velvet and a static image of a velvet fabric. This increased activity, which was observed throughout the finger exploration of the tactile device (i.e., no effect of movement direction with respect to the simulated main pile direction), could have contributed to the emergence of a velvet representation with simulation of velvet-related tactile and visual information. More specifically, the greater occipital activity could be linked to the observation made by Stilla and Sathian (2008) that visual and haptic texture-selectivity overlaps in the right occipital cortex. These authors found evidence for this bimodal texture-related selectivity in a fMRI study after contrasting shape-related selectivity and visual and haptic texture-selectivity. The LOC, which also showed greater activity in the condition with velvet-like dynamic tactile and visual stimulations, is known to be responsive to visual and tactile inputs (Amedi et al., 2001; Stilla and Sathian, 2008). However, there is clear evidence that the role of the LOC goes beyond the mere stimuli recognition, and that this functional region is part of a network involved in object representation and recognition. For instance, Kim and Zatorre (2011) showed that the LOC is commonly active during shape discrimination through different sensory modalities (visual, tactile, and auditory). Based on this finding and on the fact that the middle longitudinal fasciculus (i.e. main long-range fiber bundle) courses from the LOC through the inferior parietal lobule (Palejwala et al., 2020), we suggest that providing a visual rendering of a virtual surface explored by a finger may have prompted the velvet-like representation and perception.

Finally, we found that compared to the exploration of the virtual velvet, the exploration of the real velvet resulted in greater activation in frontal areas as the supplementary motor and the dorsal premotor cortices. Because these motor areas respond to tactile stimulation (Graziano and Gandhi, 2000; Kim et al., 2015), our results are consistent with a greater tactile stimulation intensity in the condition with real velvet than in the condition with virtual velvet. However, further experiments are required to elucidate the reasons why the different activity of the motor areas between real and virtual velvet was mainly observed when the finger moved against pile when the participants saw a static image of the velvet fabric.

## 5. Conclusion

Reproducing the feeling of texture fabrics with tactile devices is a challenging goal, which is now particularly relevant with the advances in communication technology and the increase number of e-commerce transactions. This issue was tackled in the present work using methodologies and concepts belonging to the fields of tribology, psychophysics and neurophysiology. We found that the surface of a tactile device was felt as being rougher when one could see a synthesized image of a velvet fabric (i.e., a fabrics qualified in the literature as being rough) while exploring the device with a finger. However, to enable this perceptual outcome, two key elements were necessary. First, the tactile device had to reproduce the interaction between the finger and the velvet from tribological acquisition. Second, the velvet image had to simulate the traces that would have been left if the participants were actually sliding their index finger on a real velvet. Regions of the posterior cortex that contribute to visuo-tactile integration and which were specifically activated when these two conditions were met, could have been instrumental to the enhanced rough perception. Although the tactile device was felt as being rougher, the participants did not feel that they were touching a velvet fabric. Future work will be necessary to determine if the sensation of hairy fabrics is strengthened when using tactile devices capable of simulating both the FIV and the thickness properties of these fabrics.

## Acknowledgments

We thank Professor Betty Lemaire-Semail, from L2EP/IRCICA (University of Lille), for making available the STIMTAC tactile simulator with the support of the GDR TACT (CNRS 2033). We thank Franck Buloup for developing the software Docometre used for data acquisition, Marcel Kaszap for developing the software Analyse used for data processing. We thank all the volunteers from the University of Haute Alsace.

## Funding

This work was supported by the project COMTACT (ANR 2020-CE28-0010-03), funded by the french “Agence Nationale de la Recherche” (ANR).

## Notes

### Competing Interest Statement

The authors have declared no competing interest.

